# Spontaneous and double-strand break repair-associated quasipalindrome and frameshift mutagenesis in budding yeast: role of mismatch repair

**DOI:** 10.1101/2023.08.09.552703

**Authors:** Neal Sugawara, Mason J. Towne, Susan T. Lovett, James E. Haber

## Abstract

Although gene conversion (GC) in *Saccharomyces cerevisiae* is the most error-free way to repair double-strand breaks (DSBs), the mutation rate during homologous recombination is 1000 times greater than during replication. Many mutations involve dissociating a partially-copied strand from its repair template and re-aligning with the same or another template, leading to -1 frameshifts in homonucleotide runs, quasipalindrome (QP)-associated mutations and microhomology-mediated interchromosomal template switches. We studied GC induced by HO endonuclease cleavage at *MAT*α, repaired by an *HMR::Kl-URA3* donor. We inserted into *HMR*::*Kl-URA3* an 18-bp inverted repeat where one arm had a 4-bp insertion. Most GCs yield *mat::Kl-ura3::QP+4* (Ura^-^) outcomes, but template-switching produces Ura^+^ colonies, losing the 4-bp insertion. If the QP arm without the insertion is first encountered by repair DNA polymerase and is then (mis)used as a template, the palindrome is perfected. When the QP+4 arm is encountered first, Ura^+^ derivatives only occur after second-end capture and second-strand synthesis. QP+4 mutations are suppressed by mismatch repair (MMR) proteins Msh2, Msh3, and Mlh1, but not Msh6. Deleting Rdh54 significantly reduces QP mutations only when events creating Ura^+^ occur in the context of a D-loop but not during second-strand synthesis. A similar bias is found with a proofreading-defective DNA polymerase mutation (*pol3-01*). DSB-induced mutations differed in several genetic requirements from spontaneous events. We also created a +1 frameshift in the donor, expanding a run of 4 Cs to 5 Cs. Again, Ura3^+^ recombinants markedly increased by disabling MMR, suggesting that MMR acts during GC but favors the unbroken, template strand.

## INTRODUCTION

Faithful DNA replication is essential for the survival of all organisms (Cox *et al*. 2000; Courcelle *et al*. 2004). In humans, the accumulation of mutations has profound implications for health. Increased mutability is associated with cancer proneness, immune deficiency, premature aging and neurological and developmental impairments (Cortez 2019; Brown and Freudenreich 2021; D’Amico and Vasquez 2021; Caldecott 2022). Mutations and genomic rearrangements promote cancer, affect reproductive success and can lead to human genetic diseases.

Mutations that reshape genomes derive from many sources. Many frequent mutations arise in repetitive DNA sequences, where mispairing followed by DNA synthesis generates mutations or genomic rearrangements (Lovett 2004, 2017; Anand *et al*. 2014). This class of mutagenesis, called “template-switching”, is intrinsically different in mechanism from other types that result from DNA miscoding from physical damage to the bases of the DNA template or to alterations in nucleotide pools. Genetic and biochemical analyses have defined many cellular processes that promote or deter mutagenesis by base damage; in contrast, much less is known about the cellular pathways that affect template-switch-derived mutagenesis.

Template-switching can occur in imperfect inverted repeat sequences, known as “quasipalindromes”, where one arm of the palindromic region templates synthesis from the other. Mutations at QP sites (”QPM”) are often found as hotspots of mutation in viruses, bacteria, yeast and other genetic model systems (reviewed in (Ripley 1982; Lovett 2017)). Evidence for template-switch mutations at QP sites during evolution can also be deduced from genomic sequences (Bissler 1998; Noort *et al*. 2003; Loytynoja and Goldman 2017; Abraham and Hazkani-Covo 2021; Walker *et al*. 2021; Loytynoja 2022).

To study template-switch mutagenesis at QP sites, we previously developed specific reporters in the *lacZ* gene of the bacterium *Escherichia coli* that revert to Lac^+^ by a template-switch reaction in an 18-bp quasipalindrome. These specific mutational reporter strains allowed us to deduce that DNA exonucleases ExoI and ExoVII protect *E. coli* cells from template-switching and that these mutations occur during DNA replication (Seier *et al*. 2011, 2012; Laranjo *et al*. 2017). In addition, we have defined several mutagens for QPM including DNA replication inhibitors HU (Seier et al. 2011) and azidothymidine (Seier et al. 2012), as well as DNA/protein crosslink promoting agents, including formaldehyde, 5-azacytidine and fluoroquinolone antibiotics(Laranjo *et al*. 2019; Klaric *et al*. 2021).

Here, using a similar approach, we examine QP mutagenesis (QPM) in the budding yeast *Saccharomyces cerevisiae.* We examine events arising spontaneously, presumably during DNA replication, and those that occur during the DNA synthesis that is required for the repair of a chromosomal double-strand break (DSB) by gene conversion. Budding yeast is an ideal genetic system to study eukaryotic QPM, since much is known about DNA replication and repair, and it is easy to generate large populations to measure mutation frequencies. QPM was first discovered by Sherman and colleagues (Hampsey *et al*. 1988) in which a mutational hotspot in the *S. cerevisiae CYC1* (cytochrome oxidase) gene occurred at a 7 bp QP site; therefore, this template-switching mechanism clearly operates in budding yeast.

We have developed a sensitive system to detect mutations arising during DSB-mediated gene conversion in budding yeast (Hicks *et al*. 2010). A site-specific DSB is created rapidly and synchronously by expression of galactose-inducible HO endonuclease, cleaving the *MAT*α locus (Figure 1). The DSB is repaired by copying homologous sequences at the heterochromatic *HMR* donor, located 100 kb distally on the same chromosome arm, into which is embedded a *Kluyveromyces lactis URA3* gene (*Kl-URA3*). *Kl-URA3* is silenced by the heterochromatic structure within *HMR* but is expressed when these sequences are copied into the *MAT* locus, replacing the Yα sequences with *Kl-URA3* (Fig. 2A). Nearly all cells in the population complete this gene conversion repair event in one cell cycle. Ura3^-^ mutations that arise during the switching process can be easily recovered by plating cells on 5-fluoroorotic acid (FOA). Mutations proved to be 1000 times more frequent than spontaneous mutations of *MAT::Kl-URA3* (Hicks *et al*. 2010). That these mutations arose during the repair event could be demonstrated by un-silencing *HMR*, using the Sir2 inhibitor, nicotinamide; when *HMR::Kl-URA3* is expressed the Ura3^-^ cells become Ura3^+^, since the *HMR* donor, harboring a functional *Kl- URA3* gene, is unaltered. About 60% of the mutations arising during gene conversion are single base-pair substitutions, but many of the mutants have sequence length changes and many of these appear to have involved the dissociation of the partially copied DNA strand from its template, followed by reannealing at a microhomology. Among these events are -1 (but rarely +1) frameshift (FS) changes in homonucleotide runs, intragenic deletions, and quasipalindrome-mediated events.

**Figure 1.**
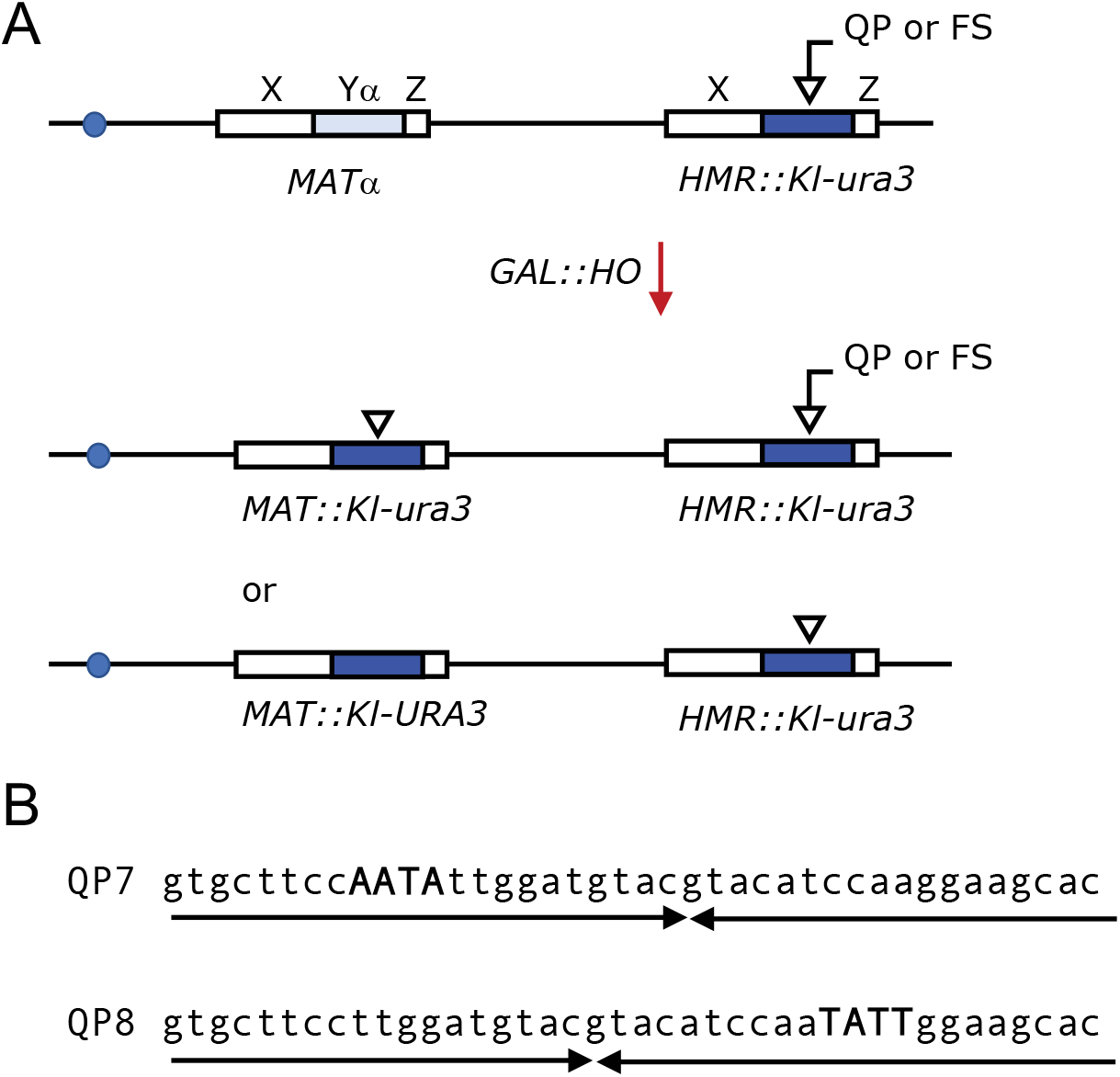
A. DSB-induced gene conversion of *MAT*α to *MAT::Kl-ura3*. Galactose-induced expression of HO endonuclease creates a DSB at *MAT*α that initiates gene conversion using *HMR::Kl-URA3* as a donor template. *Kl-URA3* replaces *HMR* Y**a**1 and possesses a quasipalindrome (QP) or frameshift (FS) mutation rendering it Ura^-^. Repair of the QP or frameshift mutations during gene conversion leads primarily to Ura-colonies in which the QP or FS was copied into *MAT*, but Ura^+^ colonies arise by template switching or frameshift mutagenesis. B. The quasipalindromes QP7 and QP8 consist of 18bp inverted repeats plus a 4bp AATA insertion (bold, upper case).

**Figure 2.**
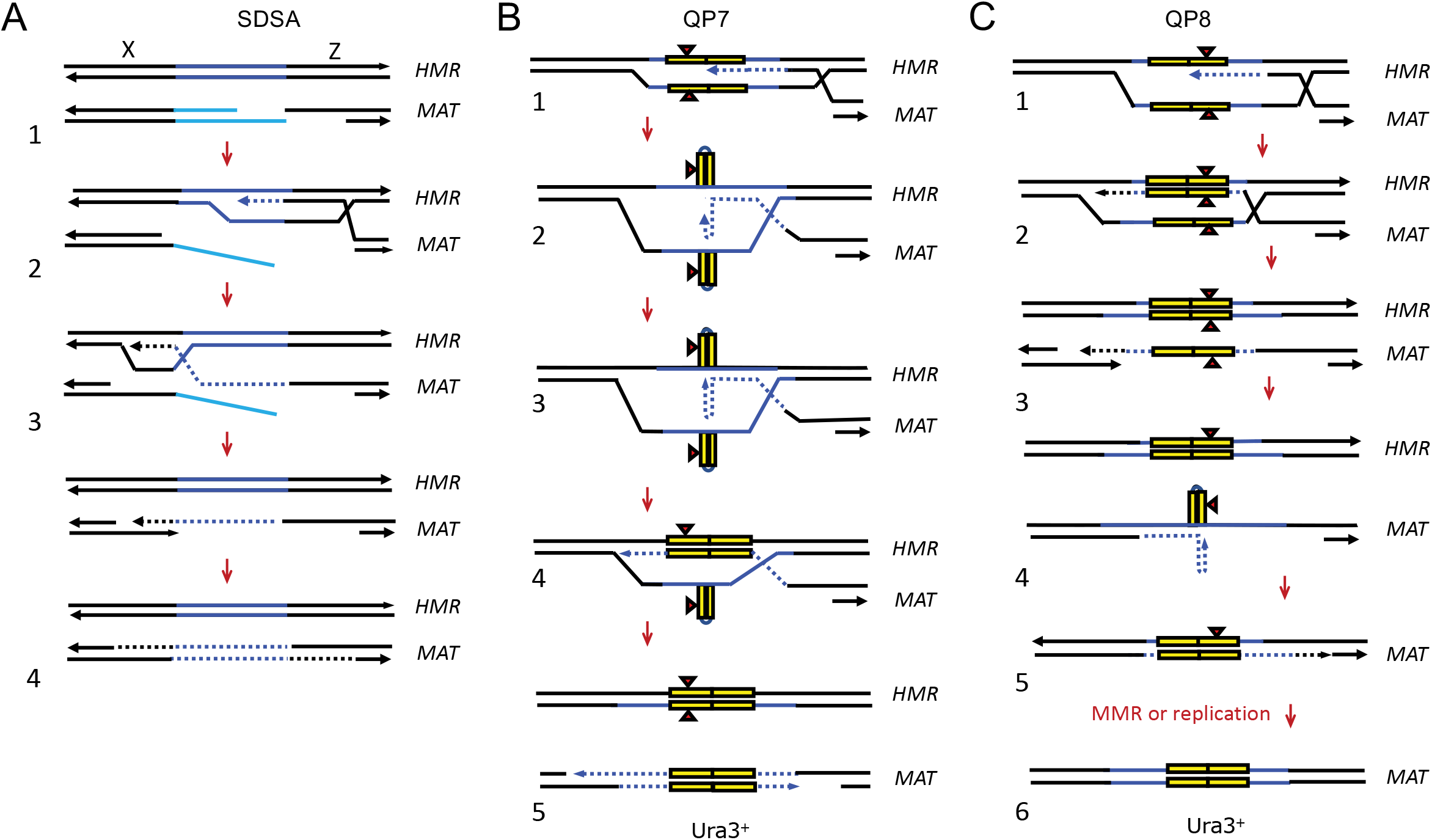
Removal of the 4 bp insertion in the quasipalindromes. **A**. In the Synthesis-Dependent Strand Annealing (SDSA) model, a DSB initiates DNA resection leaving 3’ single stranded tails. The 3’ tail strand in the Z region invades the donor sequence (HMR) forming a D-loop (1). DNA synthesis begins and the D-loop migrates to the left while displacing newly synthesized DNA (2). The displaced newly copied strand engages in second-end capture in which it anneals to the resected single stranded tail from the left side after which nonhomologous sequences at the 3’ end of the ssDNA tail are clipped off (3). The second strand of the repaired *MAT* locus is then filled in (4). **B**. In the QP7 strain the resected DNA from *MAT*-Z invades *HMR*-Z, creating a D-loop, and initiates DNA synthesis (1). As the polymerase passes through the first 18-bp repeat and enters the second repeat a hairpin forms (2). DNA synthesis continues using the first half of the QP, thus not incorporating the 4-bp insertion (red triangle) (3). The newly-synthesized strand then realigns with the donor to complete copying the donor into the *MAT*-X (4). As SDSA progresses the newly synthesized strand is displaced and anneals with the resected DNA from *MAT*-X and completes repair (5). C. In the QP8 strain the 3’ tail from *MAT*-Z strand invades *HMR*-Z and primes synthesis through QP8 (1). In the SDSA model the invading strand unwinds and is displaced from HMR (2). This allows it to anneal to the ssDNA from *MAT*-X that is formed as resection proceeds leftward from the DSB (3). Extension of this DNA to the QP sequences allows a hairpin to form and DNA synthesis to continue without incorporating the 4bp insertion (4). The final product contains a heteroduplex that can be resolved by mismatch repair or by DNA replication and mitotic segregation (5).

Quasipalindrome-mediated changes can be difficult to recognize, since they have imperfect inverted repeats that can be separated by a variable number of base pairs. Therefore, we sought to study these events by creating a specific QP that would be mutated preferentially during gene conversion. We inserted an 18-bp perfect inverted repeat, interrupted on one side or the other by a 4-base insertion (Figure 1B), thus causing the *Kl-URA3* gene to be frame-shifted and phenotypically Ura3^-^ when expressed. QP mutations arising during DSB repair that perfect the palindrome and remove the 4-bp insertion would then become Ura3^+^ (Figure 1A).

In a similar fashion we have also assayed a specific frameshift (FS) reversion event arising during DSB repair, namely the loss of one C from a homonucleotide runs of 5Cs at a “hotspot” we had previously identified in *Kl-URA3* during gene conversion repair of the HO DSB (Hicks *et al*. 2010). A +1 frameshift was made by inserting an extra C into a run of 4 Cs within the *Kl- URA3* ORF; thus Ura3^+^ cells arise if one of the Cs is deleted during gene conversion, restoring the open reading frame.

We compared Ura3+ frequencies of these HO-induced events to those obtained spontaneously, without HO cleavage. We find that there are some significant differences in the genetic requirements for QP mutatgenesis in spontaneous and DSB-induced events.

## RESULTS

### Design

To measure quasipalindrome-associated mutations accompanying gene conversion, we started with a *sir3*ti derivative of strain WH50 that can undergo gene conversion by creating a DSB at *MAT*a and using *HMR::Kl-URA3* as a donor (Hicks *et al*. 2010) The donor sequence consisted of the *Kluyveromyces lactis URA3* gene (*Kl-URA3*) inserted at *HMR* in place of the *HMR***a**1 coding sequence and removing the HO cleavage site (Fig. 1A). The *Kl-URA3* sequence was modified by Cas9-mediated gene editing to contain a QP composed of inverted 18-bp repeats plus a 4-bp insertion in either the left repeat (QP7) or the right repeat (QP8) (Figure 2). In a *sir3*Δ background, unsilencing *HMR*, these strains are Ura3^-^. Upon induction of *GAL::HO* cells repair the DSB at *MAT* by gene conversion, using *HMR:: Kl-ura3-QP* as the donor. The great majority of cells remain Ura^-^ and retain the 4-bp insertion within the QP sequence; however, in a small proportion of repair events, the 4-bp insertion is lost (i.e. the quasipalindrome is perfected) and cells become Ura3^+^. Ura3^+^ cells arose at a frequency of 6.7 or 2.1 x 10^-6^ for QP7 or QP8, respectively; the three-fold difference is statistically significant (Table 1). We note that all of the events producing Ura^+^ revertants occur during the single cell cycle in which the DSB at *MAT* is repaired by gene conversion; hence the frequencies that we observe are in fact the rates of QP mutation accompanying repair. To confirm that the Ura3^+^ recombinants were indeed corrections of the QP frameshift within the inverted repeats, we sequenced 10 independent recombination-generated Ura^+^ colonies. For both QP7 and QP8, each of the gene convertants had lost the 4-bp insertion within the QP and were therefore bona fide QP mutation events associated with gene conversion.

**Table 1.** Values for spontaneous Ura^+^ reversion obtained by the indicated mutational reporters.

| Spontaneous |  |  |  |  |  |  |
| --- | --- | --- | --- | --- | --- | --- |
| Genotype | Mean Frequency ± SEM ($\times 10^{-9}$) | | | Fold Change from WT | | |
|  | QP7 | QP8 | FS | QP7 | QP8 | FS |
| WT | 3.7 ± 0.7 | 3.4 ± 1 | 18.6 ± 9.1 | 1 | 1 | 1 |
| *msh2*Δ | 239 ± 44.7 | 110 ± 52.2 | 9410 ± 1280 | 64.6 | 32.4 | 505.9 |
| *msh3*Δ | 164 ± 63.9 | 85.3 ± 29.4 | 5940 ± 1080 | 44.3 | 25.1 | 319.4 |
| *msh6*Δ | 4.8 ± 2.6 | 1.2 ± 0.5 | 508 ± 139 | 1.3 | 0.3 | 27.3 |
| *mlh1*Δ | 509 ± 174 | 673 ± 580 | 22100 ± 4730 | 137.6 | 197.9 | 1188.2 |
| *pol2-04* | 4.9 ± 1.6 | 1.9 ± 0.7 | 8.0 ± 2.7 | 1.3 | 0.6 | 0.4 |
| *pol3-01* | 1.3 ± 0.5 | 8.5 ± 5.2 | 414 ± 55.2 | 0.4 | 2.5 | 22.3 |
| *pol32*Δ | 64 ± 58.4 | 1.2 ± 0.7 | 2.2 ± 0.9 | 17.3 | 0.4 | 0.1 |
| *rev3*Δ | 3.5 ± 1.1 | 10.2 ± 8.3 | 125 ± 110 | 0.9 | 3.0 | 6.7 |
| *rad30*Δ | 5.2 ± 1.8 | 2.1 ± 1.1 | 20.3 ± 14.4 | 1.4 | 0.6 | 1.1 |
| *rdh54*Δ | 7.1 ± 3.4 | 0.7 ± 0.2 | 9.4 ± 4.3 | 1.9 | 0.2 | 0.5 |

The frequency of Ura3^+^ events arising during homologous recombination is much higher than the spontaneous rate of QP reversion. We tested for spontaneous correction to Ura3^+^ at the *HMR::Kl-ura3-QP* sequence in *sir3*ti strains that allow gene expression of *Kl-ura3* mutants embedded within *HMR*. The frequencies of spontaneous Ura^+^ formation were assayed in strains that were deleted for *GAL::HO* since leaky expression of the endonuclease could contribute to background levels. The spontaneous frequencies were approximately 3.5 x 10^-9^ for both QP7 and QP8 (Table 1), about 1000 times lower than the HO-induced frequencies for both QP7 and QP8.

In a similar fashion we assayed a specific frameshift (FS) reversion event arising during DSB repair. In our original study we had identified a run of 4 Cs that was a hotspot for the deletion of a single C (-1C). By adding another C at this site we presumed that we would create a hotspot where the loss of one C from a homonucleotide run of now 5 Cs would restore the open reading frame. The frequency of HO-induced FS reversion (7.5 x 10^-6^) was > 300 times higher than the spontaneous frequency (Table 1). By DNA sequencing of the converted *MAT* locus in 10 independent HO-induced colonies we confirmed that each of the Ura3^+^ revertants accompanying gene conversion had indeed lost one of the 5 Cs in the FS strain.

### Effect of mismatch repair genes on QP and FS mutagenesis

The mismatch repair gene *MSH2* plays a number of roles in DSB repair (Modrich and Lahue 1996; Kolodner and Marsischky 1999; Harfe and Jinks-Robertson 2000; Oh and Myung 2022). The MutSβ heterodimer, Msh2-Msh3, acts with MutLα (Mlh1-Pms1) to recognize and correct heteroduplex DNA containing small insertion/deletions. In contrast, the Mutα (Msh2-Msh6) heterodimer, again acting with MutLα, is responsible for the mismatch correction of single base pair heterologies or insertion/deletion of single bases. The Msh2-Msh3 complex also facilitates the removal of 3’ nonhomologous flaps from DSB repair intermediates, independent of its interaction with the MutL proteins (Lyndaker and Alani 2009}. Deletion of Msh2 or Msh3 caused a > 40-fold increase in the spontaneous rate of Ura3^+^ QP mutations for both QP7 and QP8 (Figure 3) and an even larger increase (> 400-fold) in FS mutations (Figure 4). Deleting *MSH6* had little effect on either QP orientation, a result that is consistent with Msh2-Msh6 recognizing only small distortions in the DNA helix. In contrast, *msh6*ti still caused a 25-fold increase in spontaneous FS mutations. These results suggest that both Msh2-Msh3 and Msh2-Msh6 can suppress spontaneous -1 frameshifts associated with a 5-bp homonucleotide run.

**Figure 3.**
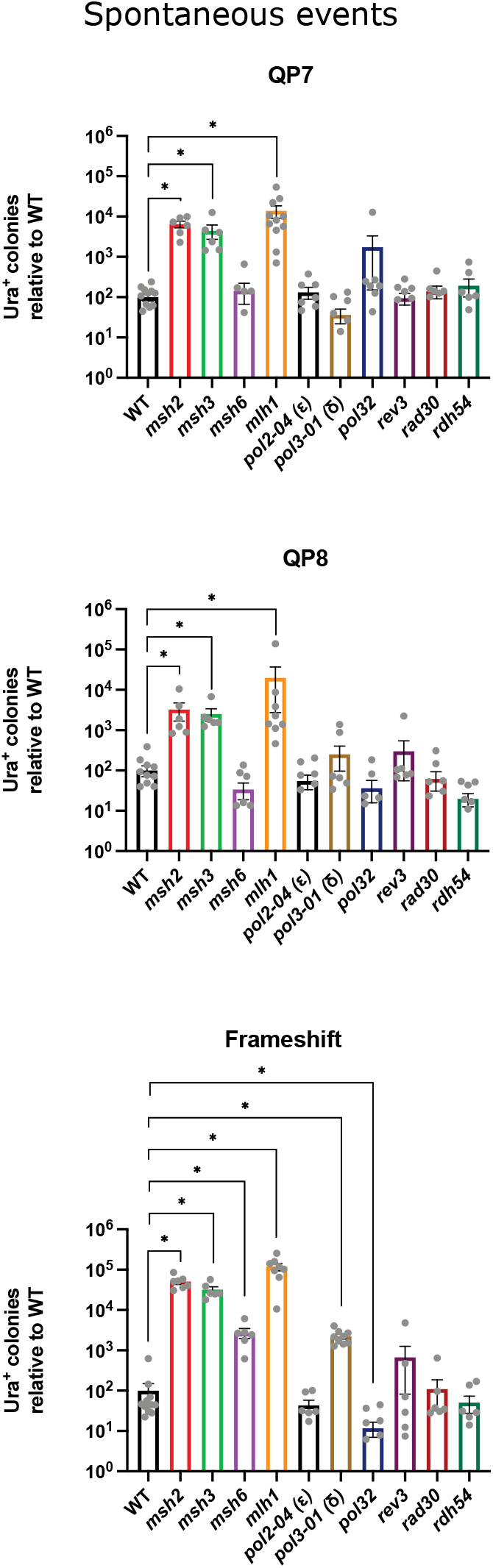
Spontaneous frameshift and QP-associated mutagenesis. Spontaneous frequencies of QP correction were measured by first integrating Kl-*URA3* at the HMR locus in a *sir4* background to allow expression. The QP7, QP8 or FS mutations were introduced to make the strains auxotrophic for uracil. Cells were cultured in rich media (YPD), appropriately diluted and plated on YPD and on selective medium lacKlng uracil to assay spontaneous correction to Ura^+^. Values were normalized to the WT frequencies (WT=100). Asterisks indicate statistical significance below p=0.005 using the Mann-Whitney test. Error bars represent the standard error of the mean.

**Figure 4.**
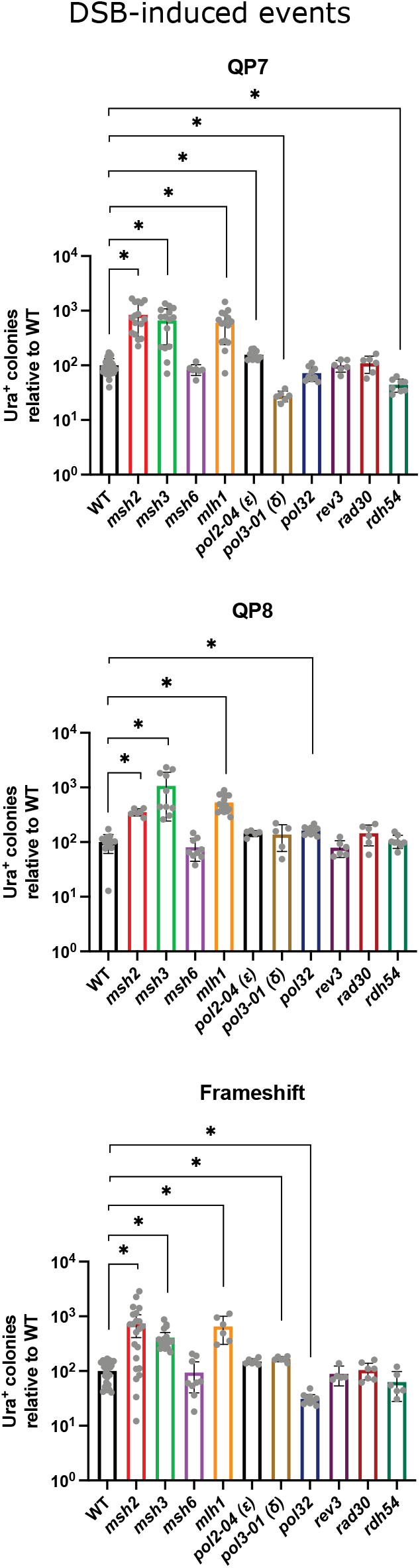
DSB-induced frameshit and QP-associated mutagenesis. Ura^+^ reversion associated with quasipalindromes, QP7, QP8, or the frameshifting during gene conversion was assayed by inducing a DSB at *MAT* using the HO endonuclease. Gal::HO was induced by plating cells on galactose-uracil medium as described in the Materials and Methods section. The rates at which QP7, QP8 or the frameshift were removed, giving rise to Ura^+^ colonies, were plotted relative to WT (WT=100). Asterisks indicate statistical significance below p=0.005 using the Mann-Whitney test. Error bars represent 95% confidence intervals.

During HO-induced DSB repair, both *msh2*ti and *msh3*ti caused a significant increase in both QP and FS mutations, though these increases (3.5 - 8 fold) were much less profound than the effects on spontaneous events (Figure 3). In contrast, *msh6*ti had no significant effect, even on the FS reversion (Figure 4). This result suggests that there are intermediates of mismatch correction during normal DNA replication that may be distinct from those in DSB repair, where Msh6 does not seem to participate in -1 deletions (see Discussion).

We confirmed that the Ura3^+^ recombinants arising during DSB repair in *msh2*ti strains were the result of the specific correction of the QP insertions or the +1 frameshift. We PCR-amplified and sequenced *MAT::KL-URA3* from 10 Ura^+^ colonies from the *msh2*ti QP7 and QP8 strains and determined that the 4-bp insertions were cleanly removed in each case. For the *msh2*ti frameshift mutation, 20 Ura3^+^ colonies were examined; 16 had removed the extra C bp from the Kl-*URA3* at *MAT*, but in 4 cases the sequence at *MAT* still had the 5C frameshift. In these 4 colonies, cells had become Ura3^+^ by a loss of one C in the *HMR::Kl-ura3-5C* locus, which in a *sir3*ti strain, is also expressed. The very high rate of spontaneous reversion of *HMR::Kl-ura3-5C* in the absence of Msh2 (Table 1) could explain these events, but the fact that the copy at *MAT* does not have this change suggests that the alteration may have happened during the switching process itself, as we had seen in a study of interchromosomal template switches (Tsaponina and Haber 2014) (see Discussion).

Mlh1-Pms1 is recruited by Msh2-Msh3 or Msh2-Msh6 and creates a nick in the DNA, in preparation for nucleolytic removal of mismatches (Modrich and Lahue 1996; Kolodner and Marsischky 1999). We examined the role of Mlh1 in the QP and the FS strains. Like *msh2*ti, *mlh1*ti mutants had a highly significant increase in the level of spontaneous mutation for QP7, QP8 and FS strain compared to the wild type (Figures 3 and 4). Among the DSB-induced events, *mlh1*Δ again behaved similarly to *msh2*ti, with a 6-fold increase in Ura3^+^ recombinants. Thus, both QP and FS correction during DSB repair require both Msh2-dependent recognition and Mlh1-dependent processing. Again, the much larger effects of *mlh1*ti on spontaneous mutations suggest differences in the way the QP and FS mutations are processed during DNA replication and in repair.

### Effect of DNA replication mutations on QP and FS mutagenesis during DSB repair

Our understanding of QP mutagenesis requires that the partly-copied DNA strand dissociate from its template in order that the end can anneal to itself, allowing the perfection of the palindrome in the newly-copied DNA but leaving the donor sequence unaltered (Figure 2). With this in mind, we surveyed strains with mutations in a number of polymerase components such as Pol2, Pol3, Pol32, Rev3, and Rad30. Pol3 is the catalytic unit of DNA polymerase δ; the *pol3-1* mutant is defective in 3’ to 5’ exonucleolytic error correction. Our previous study had shown that Pol3-01 dramatically reduced template switching events while increasing missense mutations (Hicks *et al*. 2010). An increase in single-nucleotide variants is expected in the absence of proofreading, but the decrease in template switching suggested that the mutant polymerase may dissociate less often from its template, as shown in an *in vitro* study ((Jin *et al*. 2005). Here, the *pol3-1* derivative of QP7 showed a significant 4-fold decrease relative to wild type; however, this reduction was not seen in the QP8 strain (Figure 3). The different effect of *pol3-01* may reflect the important difference in the origin of Ura3^+^ events in the two QP orientations (see Discussion). Spontaneous mutagenesis measured with QP7 was also reduced by *pol3-1*. The *pol3-01* mutation significantly increased FS reversion during DSB repair, suggesting that its defect in proofreading a -1 deletion may outweigh any effect on its processivity (Figure 4). Pol3-01 also led to a 20-fold increase in spontaneous FS reversions.

Pol2-4, a proofreading-defective mutation of DNA polymerase ε, showed a small but significant increase in frequency in the QP7 strain relative to the wild type after gene conversion (1.6x), but had no significant effect on QP8 or FS reversion (Figures 3 and 4). There were no significant effects among spontaneous events as well.

Pol32 is a nonessential subunit of Polδ that is important for break-induced replication where there is extensive DNA synthesis after strand invasion (Lydeard *et al*. 2007). In our DSB-induced assay *pol32*ti significantly decreased the number of frameshift events (3.1-fold relative to wild type) whereas it slightly increased the number of QP8 corrections (1.6x). In addition to being part of Polδ, Pol32 along with Pol31, also forms a complex with Rev3 and Rev7 as part of Polζ (Johnson *et al*. 2012; Makarova *et al*. 2012). Rev3 is the catalytic component of Polζ and is responsible for error-prone damage repair. Examination of the *rev3*ti mutant showed no significant difference from wild type indicating that the effects of *pol32*ti occur largely through its association with Polδ (Figures 3 and 4). Rad30 is DNA polymerase eta, a translesion repair polymerase. The *rad30*ti mutant showed no effect on the QP or frameshift mutations in the DSB-induced assays nor in spontaneous events.

### Effect of *rdh54*ti affecting template switching on QP and FS mutagenesis

In our previous study of DSB repair events accompanying *MAT* switching using *HMR::Kl-URA3* as the donor, we discovered that a surprisingly high number DSB-associate mutations had engaged in interchromosomal template switching (ICTS) in which the copying of *HMR::Kl-URA3* was interrupted by two template switching events, into and out of a 72% identical *S. cerevisiae ura3-52* gene on a different chromosome (Tsaponina and Haber 2014). Among deletions of genes known to be involved in homologous recombination, only one - *RDH54* - had a highly selective effect on template switching. An *rdh54*ti strain had no marked defect in simple gene conversions - replacing *MAT*α by *MAT::Kl-ura3* - but was > 50x decreased for ICTS events (Tsaponina and Haber 2014). Deleting Rdh54 also markedly reduced template jumps in a break-induced replication assay (Anand *et al*. 2014). Here we show that QP7 but not QP8 events are reduced by *rdh54*ti; also FS events were not reduced by deleting Rdh54 (Figures 3 and 4). The differences between the two QP orientations is again significant and suggests that Rdh54’s role is linked to the structure of the recombination intermediates distinguishing QP7 from QP8 (see Discussion). Spontaneous events detected with QP8 were also modestly reduced by *rdh54*ti.

## DISCUSSION

From our analysis of quasipalindrome mutagenesis and frameshift we conclude that there are significant differences between spontaneous and DSB repair-associated events, but there are also notable differences depending on the orientation of the QP relative to the direction of DNA synthesis. Previous studies from *E. coli* have shown that the rate of QPM is elevated on the leading strand of the normal replication fork because of single-strand DNA binding protein recruitment of exonucleases that abort QP mutagenesis on the lagging strand (Seier *et al*. 2011; Laranjo *et al*. 2017). In the absence of exonucleases, lagging strand QPM is more frequent (Seier *et al*. 2011; Laranjo *et al*. 2017), presumably because the opening of the DNA ahead of the DNA polymerase allows the template strand to form hairpin structures that should facilitate QPM.

One of the distinctive features of *MAT* switching is that the MAT-Z end of the DSB ends is perfectly matched with the *HMR::Kl-URA3* donor sequence, while the other end terminates in a ∼700-bp nonhomology that must be removed before second-end DNA synthesis can be accomplished (Figure 2A). Thus, in QP7 the 4-bp insertion in the hairpin is distal to the direction of first-strand repair synthesis (Figure 2B), while in QP8 the DNA polymerase encounters the QP arm carrying the +4 insertion first (Figure 2C). The polarity of strand synthesis accounts for some of the different effects of mutations that we find between QP7 and QP8. Differences in the composition of the normal replication fork and the machinery involved in DSB repair most likely accounts for differences in the dependence of spontaneous and DSB-induced QPM on various replication and repair factors. *HMR* is flanked by a known origin of replication on the left side, but we do not know the predominant direction of DNA replication across this region.

In theory, mismatch repair removing the 4-bp insertion could occur within a hairpin formed on the template strand before the region is copied during repair (Figure 2B). However, this would revert the *HMR::Kl-QP7* region, whereas we find that the donor sequence remains unchanged and the Ura^+^ sequence is found only at the recipient locus. We did find such reversion events creating *HMR::Kl-URA3* when Msh2 was deleted in the +1 FS strain, but in those instances, the sequence transferred to *MAT* still contained the 1-base insertion, suggesting that reversion of the QP occurred during the repair event itself. We previously observed such changes in the donor sequence in interchromosomal template switches (Tsaponina and Haber 2014).

In QP7, first-strand repair synthesis can form a hairpin such that the dissociation of the partially copied strand will permit the formation of the partial cruciform intermediate that will allow this strand to eliminate the insertion. This intermediate presumes that the two donor strands do not themselves reanneal but remain in a D-loop. We imagine that the D-loop is kept open by binding of single-strand binding protein complex, RPA, to the displaced donor strand. After DNA polymerase reaches the end of the palindrome, there must be a second template switch to re-align the sequences such that repair synthesis can continue into the HMR-X region that will allow it to - yet again - dissociate so that the second end can be captured and initiate second-strand synthesis (Figure 2B-4 and -5). However, before displacement and second-end capture, the newly copied strand, with its perfected palindrome, apparently anneals to its template strand, creating heteroduplex DNA that can be recognized and mismatch repaired by the MutSα-MutLα complex. We surmise that mismatch repair is highly directional, favoring the complete template strand over the elongating repair strand (Figure 5). This directionality of MMR will restore the 4-bp insertion on the newly copied strand and eliminate Ura3^+^ outcomes. Hence we find a very significant increase in Ura^+^ recombinants when Msh2, Msh3 or Mlh1 are deleted. Because Msh6 does not recognize 4-bp insertions, the rate of Ura^+^ in *msh6*ti is not elevated relative to the wild type control.

**Figure 5.**
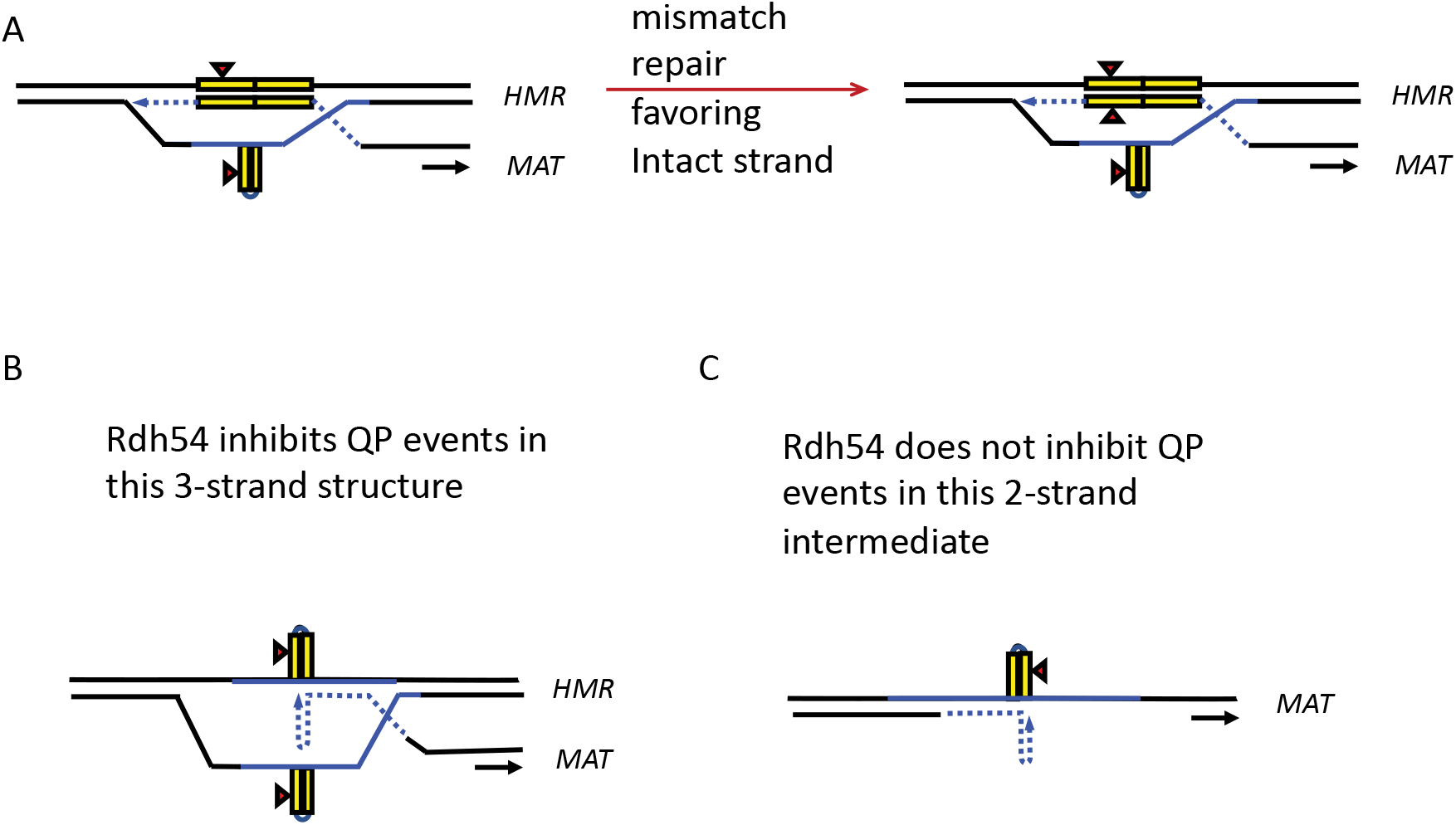
Rdh54 is important for mismatch repair of mutations in QP7 but not QP8. A D-loop containing a heteroduplexed DNA intermediate is preferentially repaired in favor of the unbroken template strand (A). In the absence of Rdh54 QP mutations in the context of a D-loop (detected by the QP7 reporter) are less likely to occur (B). Replication after strand displacement and annealingthat gives rise to QP associated mutations (detected by QP8) is not affected by Rdh54, presumably because it occurs in the ssDNA gap rather than a D-loop (C).

Although *msh2*ti, *msh3*ti and *mIh1*ti also elevate the rate of Ura^+^ events in QP8, the rates of these events and of the wildtype are all significantly lower than for QP7. This difference reflects the fact that first-strand repair synthesis cannot produce a corrected Ura3^+^ strand (Figure 3C-2), as the formation of a transient partial cruciform and copying itself will lead to the insertion of a second 4-bp segment. Thus, creating a Ura3^+^ sequence with QP8 can only occur during second-strand synthesis, when the template is not the *HMR* donor locus, but the first-strand copy that is captured by the second end of the DSB (Figure 3C-4). Dissociation of the partially-copied second strand and realignment on the first-strand template will create a heteroduplex that again can be recognized by Msh2-Msh3, and not Msh6 (Figure 3C-5). Again, MMR is likely to favor correction of the heteroduplex in favor of the “intact” strand and hence Ura^+^ events will be eliminated. In the absence of mismatch repair, the heteroduplex will persist and cells will only become Ura^+^ after a round to DNA replication that will create a Ura^+^/Ura^-^ sectored colony. Because the rate of QP events is low, it has not been possible to identify sectored colonies directly.

The much larger effect of *msh2*ti, *msh3*ti and *mlh1*ti on spontaneous frequencies relative to those induced during *MAT* switching may arise from limits to mismatch discrimination during DSB repair. During first-strand synthesis, capacity for removal of premutations exists only when they are paired in heteroduplex with their templates and because of D-loop migration, the newly replicated strand may be extruded as a single-strand shortly after synthesis, and so lacks a partner for mismatch repair. During second-strand synthesis, it may be impossible to discriminate mutant vs parental information. During normal semiconservative synthesis, there is likely a larger window to discriminate and remove mismatched nucleotides from the newly replicated strand than what occurs during DSB repair.

One striKlng difference between DSB repair-induced mutagenesis of QP7 and QP8 is the effect of *rdh54*ti. We suggest that the lack of inhibition in QP8 reflects the different DNA intermediates where mutagenesis is proposed to occur. In QP7, the dissociation of the partially copied strand takes place in the context of a migrating D-loop, whose length and stability are affected by Rdh54 (Petukhova *et al*. 2000; Piazza *et al*. 2019; Shah *et al*. 2020; Keymakh *et al*. 2022). In QP8, mutagenesis appears to occur in a simpler context, where the first-strand ssDNA has been captured by the second end, which then uses the first strand as a template; this occurs in the absence of a competing complementary strand that is present in the migrating D-loop. The different effects of deleting Rdh54 in these two contexts supports these models of mutagenesis.

We offer a similar explanation for the different effect of *pol3-01* in QP7 and QP8: in the context of a migrating D-loop the greater processivity of *pol3-01* will lead to a lower rate of new strand dissociation, whereas in QP8, *pol3-01* apparently does not change the rate of dissociation. Another explanation for differential Pol effects on QP mutations may be caused by the superior ability of Pol δ over Pol ε to catalyze strand displacement synthesis (Burgers and Kunkel 2017). 3’ extension DNA synthesis from an internally paired QP structure may require the displacement of the parental strand template, a reaction more readily catalyzed by Pol δ in vitro (Burgers and Kunkel 2017). The nonessential Pol32 subunit of Pol has been proposed to promote strand displacement in vivo (Budd *et al*. 2006} and to make Pol δ more processive in vivo.(Garbacz *et al*. 2018) For spontaneous QP mutagenesis detected with either QP7 and QP8, there are indications of effects by Pol32, Rdh54 and proofreading mutants in Pol but further work will be required to substantiate this.

In frameshift (-1C) mutagenesis, we again find differences in the roles of various repair proteins in spontaneous and repair-associated events. Here, both Msh3 and Msh6 appear to contribute to the suppression of spontaneous events, presumably along with Msh2, and Mlh1 is also implicated. In both situations, *pol3-01* leads to a significant increase in mutations, attributable to its defect in proofreading, but *pol2-04* does not.

## METHODS

### Strains

All strains are derivatives of WH050 (*ho hml*ti::*ADE1 MAT*α *hmr::Kl-URA3 ade1 leu2 Iys5 trp1::hisG ura3-52 ade3::GAL::HO*) that was altered by introducing *sir3*ti::HPHMX (TOY7). The QP consists of 18bp inverted repeats (GTGCTTCCTTGGATGTAC) with a 4bp insertion (GTGCTTCC**AATA**TTGGATGTAC) in either the left (QP7) or right (QP8) repeat. The QPs replace the wild type sequence CTCTTGACGTTCGTTCGACTGATGAGCTAT in the *Kl-URA3* open reading frame. The frameshift mutation consists of a C inserted into a run of 4 Cs at the start of the sequence CCCCAGGTGTAGGTTTAGAC in the coding sequence. Mutations were introduced by repairing a Cas9-induced DSB within *Kl-URA3* (Anand *et al*. 2017).The DSB was repaired using a synthetic 500bp dsDNA containing the pertinent sequence alterations. Deletion mutations were introduced using the KANMX drug cassette (*msh2*ti, *msh3*ti, *msh6*ti, *pol32*ti, *rdh54*ti, *rev3*ti and *rad30*ti) or the NATMX cassette (*mlh1*ti). The *pol3-1* and *pol2-4* alleles were created using Cas9 to create a DSB within *POL3* or *POL2* that was repaired by using a ssDNA oligonucleotide containing the requisite sequence changes (Gallagher *et al*. 2010). We also determined the frequencies of spontaneous mutations using derivatives deleted for *GAL::HO*. These strains were constructed by transforming the strains with a PCR product from *ADE3* that converted *ade3::GAL::HO* to *ADE3*.

### Assays

To assay spontaneous events, single colonies grown on plates containing 5-FOA were inoculated into YEPD media and grown for ∼18 hours or until saturation. Cultures were then diluted 1:10 with fresh media and incubated for 2 hours to resume growth. Cultures were then washed twice with sterile water by centrifugation then concentrated in sterile water. A fraction of the resulting resuspension was serially diluted and spotted onto YEPD plates to quantify the total cells in the sample. The remaining sample was plated on media plates lacKlng uracil. YEPD plate colonies were counted after 2 days and uracil-lacKlng plate colonies after 4 days. The frequencies of reversion events were then calculated. For strains with particularly low reversion frequencies, some assays were performed with 15mL initial cultures. The entire culture was collected after ∼18 hours of growth and then assayed as described, proceeding directly to the wash step.

For assaying DSB-induced gene conversion events, cells were grown in 5-fluororotic acid (5-FOA) to minimize the growth of spontaneous mutants, which was especially a problem in strains lacKlng mismatch repair genes that have an elevated level of spontaneous mutations. Strains were grown in 5-FOA for 2 days, washed twice with sterile water by centrifugation, briefly sonicated to break up clumps of cells, diluted and plated onto 1) selective medium lacKlng uracil and containing galactose and 2) rich medium (YPD). Colonies were counted after 3 to 5 days and the frequencies were determined. Cells were also plated on dextrose-containing plates lacKlng uracil to assess background levels of mutation correction. When rich medium was used in place of 5-FOA spontaneous events occurred at a significant level in certain strains in the *msh2*ti *msh3*ti or *mlh1*ti backgrounds which may be due to leaky repression of the *GAL::HO* gene.

When gene conversion was induced with *GAL::HO* we observed that the level of Ura^+^ events in both the QP strains and in the frameshift strain was higher than the spontaneous levels (compare Tables 1 and 2). In this assay, however, we observed a significant level of background events that were measured by plating cells on Ura^-^ plates lacKlng galactose. These events are likely due to leaky repression of the *GAL::HO* gene. In the wild type and in most of the mutants this background level was low relative to the induced levels. These ranged from 0.14% to 9.3% relative to the induced levels. The exceptions were the frameshift mutants in the *msh2*ti, *msh3*ti and *mlh1*ti backgrounds which had background levels that were respectively 32%, 11% and 42% when compared to the induced levels. For the values reported in Table 2 the background levels were subtracted from the induced levels.

**Table 2.**
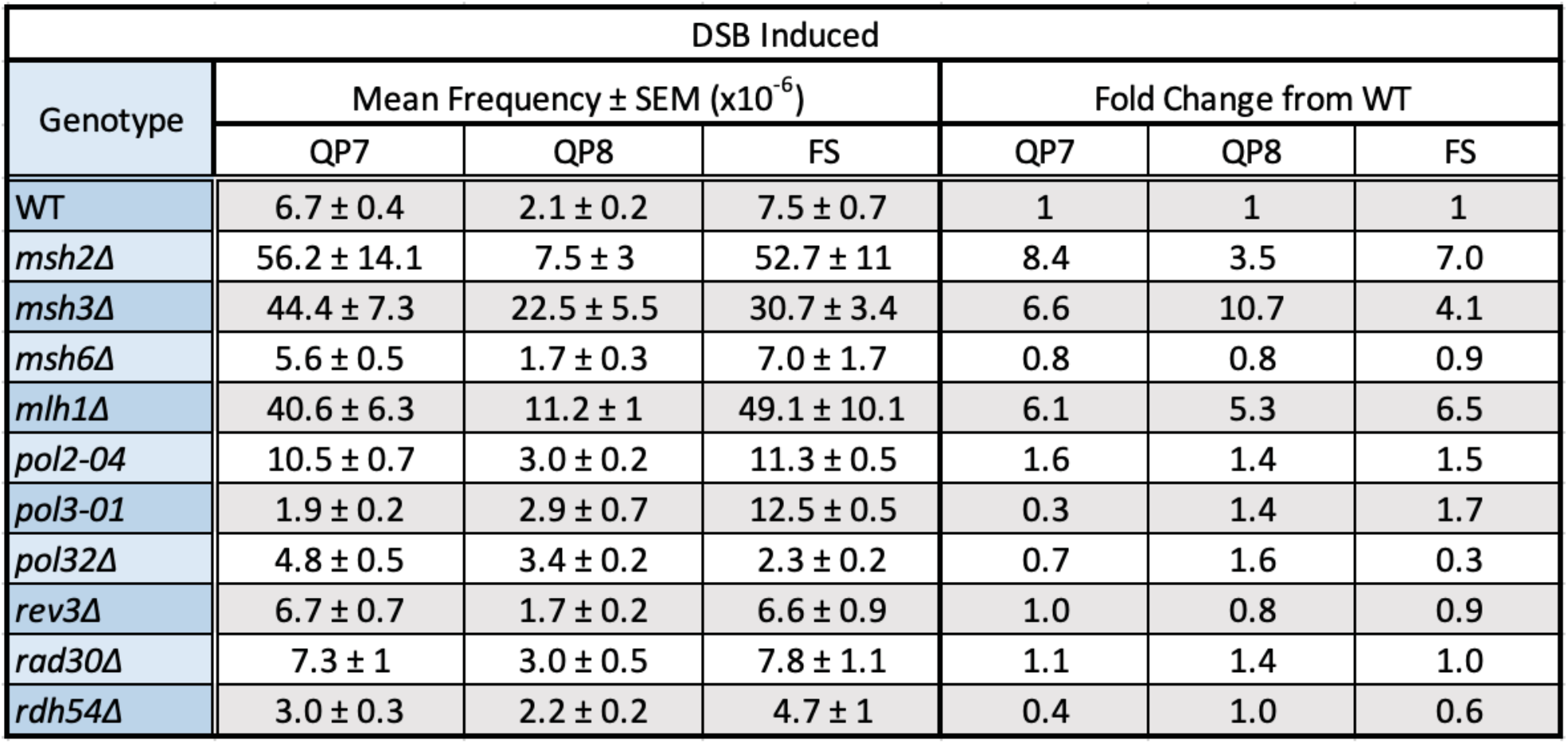
Values for DSB-induced Ura^+^ reversion obtained by the indicated mutational reporters.

| DSB Induced |  |  |  |  |  |  |
| --- | --- | --- | --- | --- | --- | --- |
| Genotype | Mean Frequency ± SEM ($\times 10^{-6}$) | | | Fold Change from WT | | |
|  | QP7 | QP8 | FS | QP7 | QP8 | FS |
| WT | 6.7 ± 0.4 | 2.1 ± 0.2 | 7.5 ± 0.7 | 1 | 1 | 1 |
| *msh2*Δ | 56.2 ± 14.1 | 7.5 ± 3 | 52.7 ± 11 | 8.4 | 3.5 | 7.0 |
| *msh3*Δ | 44.4 ± 7.3 | 22.5 ± 5.5 | 30.7 ± 3.4 | 6.6 | 10.7 | 4.1 |
| *msh6*Δ | 5.6 ± 0.5 | 1.7 ± 0.3 | 7.0 ± 1.7 | 0.8 | 0.8 | 0.9 |
| *mlh1*Δ | 40.6 ± 6.3 | 11.2 ± 1 | 49.1 ± 10.1 | 6.1 | 5.3 | 6.5 |
| *pol2-04* | 10.5 ± 0.7 | 3.0 ± 0.2 | 11.3 ± 0.5 | 1.6 | 1.4 | 1.5 |
| *pol3-01* | 1.9 ± 0.2 | 2.9 ± 0.7 | 12.5 ± 0.5 | 0.3 | 1.4 | 1.7 |
| *pol32*Δ | 4.8 ± 0.5 | 3.4 ± 0.2 | 2.3 ± 0.2 | 0.7 | 1.6 | 0.3 |
| *rev3*Δ | 6.7 ± 0.7 | 1.7 ± 0.2 | 6.6 ± 0.9 | 1.0 | 0.8 | 0.9 |
| *rad30*Δ | 7.3 ± 1 | 3.0 ± 0.5 | 7.8 ± 1.1 | 1.1 | 1.4 | 1.0 |
| *rdh54*Δ | 3.0 ± 0.3 | 2.2 ± 0.2 | 4.7 ± 1 | 0.4 | 1.0 | 0.6 |

### Sequencing

The removal of the +4 bp insertion in QPs or one C from FS at *mat::Kl-URA3* was confirmed by PCR amplifying *mat::Kl-URA3* using the primers from MAT X (TGTTACACTCTCTGGTAACTTAGG) and MAT-distal sequences (CCGCATGGGCAGTTTACCT) and sequencing the PCR products.

Strains and plasmids are available upon request. The authors affirm that all data necessary for confirming the conclusions of the article are present within the article, figures, and tables.

## BIBLIOGRAPHY

Abraham M., and E. Hazkani-Covo, 2021 Protein innovation through template switching in the Saccharomyces cerevisiae lineage. Sci Rep-uk 11: 22558. https://doi.org/10.1038/s41598-021-01736-y

Anand R. P., O. Tsaponina, P. W. Greenwell, C.-S. Lee, W. Du, et al., 2014 Chromosome rearrangements via template switching between diverged repeated sequences. Gene Dev 28: 2394–406. https://doi.org/10.1101/gad.250258.114

Anand R., G. Memisoglu, and J. Haber, 2017 Cas9-mediated gene editing in Saccharomyces cerevisiae. Protoc. Exch. https://doi.org/10.1038/protex.2017.021a

Bissler J. J., 1998 DNA inverted repeats and human disease. Frontiers in bioscience : a journal and virtual library 3: d408–18.

Brown R. E., and C. H. Freudenreich, 2021 Structure-forming repeats and their impact on genome stability. Curr 0pin Genet Dev 67: 41–51. https://doi.org/10.1016/j.gde.2020.10.006

Budd M. E., C. C. Reis, S. Smith, K. Myung, and J. L. Campbell, 2006 Evidence Suggesting that Pif1 Helicase Functions in DNA Replication with the Dna2 Helicase/Nuclease and DNA Polymerase 8. Mol Cell Biol 26: 2490–2500. https://doi.org/10.1128/mcb.26.7.2490-2500.2006

Burgers P. M. J., and T. A. Kunkel, 2017 Eukaryotic DNA Replication Fork. Annual review of biochemistry 86: 417–438. https://doi.org/10.1146/annurev-biochem-061516-044709

Caldecott K. W., 2022 DNA single-strand break repair and human genetic disease. Trends Cell Biol 32: 733–745. https://doi.org/10.1016/j.tcb.2022.04.010

Cortez D., 2019 Replication-Coupled DNA Repair. Mol Cell 74: 866–876. https://doi.org/10.1016/j.molcel.2019.04.027

Courcelle J., J. J. Belle, and C. T. Courcelle, 2004 When replication travels on damaged templates: bumps and blocks in the road. Research in microbiology 155: 231–237. https://doi.org/10.1016/j.resmic.2004.01.018

Cox M. M., M. F. Goodman, K. N. Kreuzer, D. J. Sherratt, S. J. Sandler, et al., 2000 The importance of repairing stalled replication forks. Nature 404: 37–41. https://doi.org/10.1038/35003501

D’Amico A. M., and K. M. Vasquez, 2021 The multifaceted roles of DNA repair and replication proteins in aging and obesity. Dna Repair 99: 103049. https://doi.org/10.1016/j.dnarep.2021.103049

Garbacz M. A., S. A. Lujan, A. B. Burkholder, P. B. Cox, Q. Wu, et al., 2018 Evidence that DNA polymerase 8 contributes to initiating leading strand DNA replication in Saccharomyces cerevisiae. Nat. Commun. 9: 858. https://doi.org/10.1038/s41467-018-03270-4

Hampsey D. M., J. F. Ernst, J. W. Stewart, and F. Sherman, 1988 Multiple base-pair mutations in yeast. Journal of Molecular Biology 201: 471–486. https://doi.org/10.1016/0022-2836(88)90629-8

Harfe B. D., and S. Jinks-Robertson, 2000 Mismatch repair proteins and mitotic genome stability. Mutation research 451: 151–167.

Hicks W. M., M. Klm, and J. E. Haber, 2010 Increased mutagenesis and unique mutation signature associated with mitotic gene conversion. Science (New York, NY) 329: 82–85. https://doi.org/10.1126/science.1191125

Jin Y. H., P. Garg, C. M. W. Stith, H. Al-Refai, J. F. Sterling, et al., 2005 The Multiple Biological Roles of the 3’→5’ Exonuclease of Saccharomyces cerevisiae DNA Polymerase 8 Require Switching between the Polymerase and Exonuclease Domains. Mol Cell Biol 25: 461–471. https://doi.org/10.1128/mcb.25.1.461-471.2005

Johnson R. E., L. Prakash, and S. Prakash, 2012 Pol31 and Pol32 subunits of yeast DNA polymerase 8 are also essential subunits of DNA polymerase δ. Proc National Acad Sci 109: 12455–12460. https://doi.org/10.1073/pnas.1206052109

Keymakh M., J. Dau, J. Hu, B. Ferlez, M. Lisby, et al., 2022 Rdh54 stabilizes Rad51 at displacement loop intermediates to regulate genetic exchange between chromosomes. Plos Genet 18: e1010412. https://doi.org/10.1371/journal.pgen.1010412

Klaric J. A., D. J. Glass, E. L. Perr, A. D. Reuven, M. J. Towne, et al., 2021 DNA damage-signaling, homologous recombination and genetic mutation induced by 5-azacytidine and DNA-protein crosslinks in Escherichia coli. Mutat Res Fundam Mol Mech Mutagen 822: 111742. https://doi.org/10.1016/j.mrfmmm.2021.111742

Kolodner R. D., and G. T. Marsischky, 1999 Eukaryotic DNA mismatch repair. Curr Opin Genet Dev 9: 89–96. https://doi.org/10.1016/s0959-437x(99)80013-6

Laranjo L. T., S. J. Gross, D. M. Zeiger, and S. T. Lovett, 2017 SSB recruitment of Exonuclease I aborts template-switching in Escherichia coli. DNA Repair 57: 12–16. https://doi.org/10.1016/j.dnarep.2017.05.007

Laranjo L. T., J. A. Klaric, L. R. Pearlman, and S. T. Lovett, 2019 Stimulation of Replication Template-Switching by DNA-Protein Crosslinks. Genes 10: 14. https://doi.org/10.3390/genes10010014

Lovett S. T., 2004 Encoded errors: mutations and rearrangements mediated by misalignment at repetitive DNA sequences. Molecular Microbiology 52: 1243–1253. https://doi.org/10.1111/j.1365-2958.2004.04076.x

Lovett S. T., 2017 Template-switching during replication fork repair in bacteria. DNA Repair 1–0. https://doi.org/10.1016/j.dnarep.2017.06.014

Loytynoja A., and N. Goldman, 2017 Short template switch events explain mutation clusters in the human genome. Genome Res 27: 1039–1049. https://doi.org/10.1101/gr.214973.116

Loytynoja A., 2022 Thousands of human mutation clusters are explained by short-range template switching. Genome Res 32: 1437–1447. https://doi.org/10.1101/gr.276478.121

Lydeard J. R., S. Jain, M. Yamaguchi, and J. E. Haber, 2007 Break-induced replication and telomerase-independent telomere maintenance require Pol32. 448: 820–823. https://doi.org/10.1038/nature06047

Lyndaker A. M., and E. Alani, 2009 A tale of tails: insights into the coordination of 3’ end processing during homologous recombination. BioEssays : news and reviews in molecular, cellular and developmental biology 31: 315–321. https://doi.org/10.1002/bies.200800195

Makarova A. V., J. L. Stodola, and P. M. Burgers, 2012 A four-subunit DNA polymerase δ complex containing Pol 8 accessory subunits is essential for PCNA-mediated mutagenesis. Nucleic Acids Res 40: 11618–11626. https://doi.org/10.1093/nar/gks948

Modrich P., and R. Lahue, 1996 Mismatch repair in replication fidelity, genetic recombination, and cancer biology. Annual review of biochemistry 65: 101–133. https://doi.org/10.1146/annurev.bi.65.070196.000533

Noort V. van, P. Worning, D. W. Ussery, W. A. Rosche, and R. R. Sinden, 2003 Strand misalignments lead to quasipalindrome correction. Trends Genet 19: 365–369. https://doi.org/10.1016/s0168-9525(03)00136-7

Oh J.-M., and K. Myung, 2022 Crosstalk between different DNA repair pathways for DNA double strand break repairs. Mutat Res Genetic Toxicol Environ Mutagen 873: 503438. https://doi.org/10.1016/j.mrgentox.2021.503438

Petukhova G., P. Sung, and H. Klein, 2000 Promotion of Rad51-dependent D-loop formation by yeast recombination factor Rdh54/Tid1. Gene Dev 14: 2206–2215. https://doi.org/10.1101/gad.826100

Piazza A., S. S. Shah, W. D. Wright, S. K. Gore, R. Koszul, et al., 2019 Dynamic Processing of Displacement Loops during Recombinational DNA Repair. Mol Cell 73: 1255–1266.e4. https://doi.org/10.1016/j.molcel.2019.01.005

Ripley L., 1982 Model for the participation of quasi-palindromic DNA sequences in frameshift mutation. Proc Natl Acad Sci U S A 79: 4128–4132.

Seier T., D. R. Padgett, G. Zilberberg, V. A. J. Sutera, N. Toha, et al., 2011 Insights into mutagenesis using Escherichia coli chromosomal lacZ strains that enable detection of a wide spectrum of mutational events. Genetics 188: 247–262. https://doi.org/10.1534/genetics.111.127746

Seier T., G. Zilberberg, D. M. Zeiger, and S. T. Lovett, 2012 Azidothymidine and other chain terminators are mutagenic for template-switch-generated genetic mutations. Proc Natl Acad Sci U S A 109: 6171–6174. https://doi.org/10.1073/pnas.1116160109

Shah S. S., S. Hartono, A. Piazza, V. Som, W. Wright, et al., 2020 Rdh54/Tid1 inhibits Rad51-Rad54-mediated D-loop formation and limits D-loop length. Elife 9: e59112. https://doi.org/10.7554/elife.59112

Tsaponina O., and J. E. Haber, 2014 Frequent Interchromosomal Template Switches during Gene Conversion in S. cerevisiae. Mol Cell 55: 615–625. https://doi.org/10.1016/j.molcel.2014.06.025

Walker C. R., A. Scally, N. D. Maio, and N. Goldman, 2021 Short-range template switching in great ape genomes explored using pair hidden Markov models. Plos Genet 17: e1009221. https://doi.org/10.1371/journal.pgen.1009221

